# ORF6 protein of SARS-CoV-2 inhibits TRIM25 mediated RIG-I ubiquitination to mitigate type I IFN induction

**DOI:** 10.1101/2022.05.05.490850

**Authors:** Oyahida Khatun, Mansi Sharma, Rohan Narayan, Shashank Tripathi

**Affiliations:** Emerging Viral Pathogens Laboratory, Centre for Infectious Disease Research, Indian Institute of Science, Bengaluru, 560012, India; Microbiology & Cell Biology Department, Biological Sciences Division, Indian Institute of Science, Bengaluru, 560012, India

**Keywords:** SARS-CoV-2, COVID-19, Innate Immunity, Type-I IFN, ORF6, RIG-I, Ubiquitination, TRIM25

## Abstract

Evasion and antagonism of host cellular immunity upon SARS-CoV-2 infection confers a profound replication advantage on the virus and contributes to COVID-19 pathogenesis. We explored the ability of different SARS-CoV-2 proteins to antagonize the host innate immune system and found that the ORF6 protein mitigated type-I IFN (interferon) induction and downstream IFN signaling. Our findings also corroborated previous reports that ORF6 blocks the nuclear import of IRF3 and STAT1 to inhibit IFN induction and signaling. Here we show that ORF6 directly interacts with RIG-I and blocks downstream type-I IFN induction and signaling by inhibiting K-63 linked ubiquitination of RIG-I by the E3 Ligase TRIM25. This involves ORF6-mediated targeting of TRIM25 for degradation, also observed during SARS-CoV-2 infection. The type-I IFN antagonistic activity of ORF6 was mapped to its C-terminal cytoplasmic tail, specifically to amino acid residues 52-61. Overall, we provide new insights into how the SARS-CoV-2 ORF6 protein inhibits type I-IFN induction and signaling through distinct mechanisms.

## Introduction

While the COVID-19 pandemic continues for the third year, its aetiological agent SARS-CoV-2 stays in the focus of intense scientific investigation by researchers worldwide. Tremendous progress has been made on the front of vaccine and antiviral development against COVID-19, but the basic biology of the virus is still being explored. SARS-CoV-2 is a single-stranded positive-sense RNA virus of the family *Coronaviridae* that harbors at least 4 known seasonal coronaviruses and more pathogenic SARS and MERS coronaviruses (Coronaviridae Study Group of the International Committee on Taxonomy of, 2020). The genome comprises two overlapping open reading frames (ORFs), ORF1a and ORF1b, which are translated to generate continuous polypeptides and subsequently cleaved into 16 non-structural proteins (NSPs) (Finkel *et al*, 2021). One shared aspect of *Betacoronaviruses* is the ability to evade and antagonize the host’s innate and adaptive immune responses (Li *et al*, 2021). This property is essential for efficient virus infection and replication, contributing to viral pathogenesis. Especially in the case of COVID-19, dampening of early cellular innate immune response and subsequent dysregulation of cytokine expression is considered a significant contributor to severe COVID-19. Hence, a detailed investigation into viral mechanisms of host immune evasion and antagonism is essential for developing effective therapeutic interventions.

One of the earliest cellular antiviral responses is orchestrated by IFNs, which pose a crucial hurdle that viruses need to overcome upon infection. IFN response begins with recognition of viral Pathogen-associated molecular patterns (PAMPs) by cellular Pattern Recognition Receptors (PRRs), which relay the message through specific kinases to transcription factors IRF3, IRF7, and NF□B, which in turn induce expression of type I, II or III IFNs (Mohammad *et al*, 1986), (Platanias, 2005), (Hiscott *et al*, 2006). Subsequently, IFNs activate the JAK-STAT pathway in an autocrine or paracrine manner, leading to the expression of a battery of interferon-stimulated genes (ISGs) that act as viral restriction factors and regulators of innate and adaptive immunity (Darnell *et al*, 1994). Viruses have evolved a plethora of mechanisms to inhibit IFN induction and subsequent signaling events to dampen host immunity (Garcia-Sastre, 2017), and SARS-CoV-2 is no exception (Lei *et al*, 2020), (Xia *et al*, 2020). Many SARS-CoV-2 accessory and NSPs have antagonistic effects on IFN responses by directly targeting viral sensors or blocking the downstream antiviral signaling molecules (Li *et al*., 2021)).

In this study, we screened the IFN-antagonistic ability of SARS-CoV-2 proteins and found ORF6, among others, to be the most potent inhibitor of both IFN induction and signaling. We mapped these activities to the cytoplasmic tail of ORF6, specifically the residues 52-61, which are highly conserved. Our data were coherent with earlier studies where ORF6 was shown to inhibit IFN response by blocking the nuclear import of key transcription factors involved in IFN response. However, these are downstream events, and the molecular basis of the highly efficient shutdown of IFN induction by SARS-CoV-2 ORF6 protein was not clear. We show that ORF6 directly interacts with RIG-I and blocks its ubiquitination by E3 ligase TRIM25. More specifically, the K63-linked ubiquitination of RIG-I, required for its stability and activation, is blocked by SARS-CoV-2 ORF6 protein. SARS-CoV-2 infection and ORF6 expression alone lead to TRIM25 degradation, which was reversed upon inhibition of proteasomes and autophagy. Overall, our data shed information on the molecular mechanisms of IFN-antagonism mediated by SARS-CoV-2 ORF-6 protein.

## Results & Discussion

### Multiple SARS-CoV-2 proteins antagonize Type-I IFN induction and signaling

SARS-CoV-2 has a 29.7 kb genome, 2/3^rd^ of which from the 5’ end encodes ORF1a/1b, which in turn produces 16 NSPs after proteolytic processing of polyprotein 1a and 1ab (pp1a and pp1ab); the 3’ end of the genome encodes multiple sub-genomic RNAs that encode 4 structural proteins [Spike (S); Envelope (E); Membrane (M); and Nucleocapsid (N)] and at least 9 accessory proteins (ORF3a; 3b; 6; 7a; 7b; 8; 9b; 9c and 10) (Wu *et al*, 2020a), (V’Kovski *et al*, 2021). To identify the SARS-CoV-2 proteins which may interfere with type-I IFN induction, plasmids expressing SARS-CoV-2 proteins were co-transfected with an interferon-beta (IFNβ) promoter-driven Firefly luciferase reporter plasmid (pIFNβ-FFLuc), 3), and a control plasmid constitutively expressing renilla luciferase gene (pRL-TK). The cells were treated with poly (I:C) to stimulate the type-I IFN induction pathway, and relative luciferase units were calculated. We observed that 4 proteins, namely NSP1, NSP13, NSP14, and ORF6, reduced IFN reporter induction to less than 25% of control (Fig 1A). Similarly, to identify the SARS-CoV-2 proteins which may interfere with type-I IFN signaling and ISG induction, we co-transfected SARS-CoV-2 plasmids with an ISRE promoter-driven Firefly luciferase reporter plasmid (pIFNβ-FFLuc), 3), along with control plasmid constitutively expressing renilla luciferase gene (pGL4). Cells were treated with universal interferon to stimulate type I-IFN-signalling and ISG induction. Relative Luciferase Units (RLUs) were calculated, and as before, we observed that the previous 4 proteins inhibited ISG induction to less than 25% of control (Fig 1B). We then performed western blotting to ensure that these effects on IFN induction and signaling were consistent with SARS-CoV-2 protein expression. While all constructs were expressed at variable levels, the NSP11, Orf3b, and Orf7b constructs did not produce a detectable band by western blot (Sup Figure 1A). Apart from the earlier 4 proteins, a few additional proteins also inhibited IFN induction (NSP5) or signaling (NSP7, NSP9, ORF9b), though less effectively (Sup Fig 1B). To further substantiate our results, we compiled and compared the data from previous studies where similar reporter-based approaches were utilized to screen for IFN antagonists of SARS-CoV-2 (Fig 1C, D) (Lei *et al*., 2020), (Li *et al*, 2020), (Xia *et al*., 2020), (Yuen *et al*, 2020), (Zhang *et al*, 2021), (Hayn *et al*, 2021), (Stukalov *et al*, 2021), (Shemesh *et al*, 2021), (Fu *et al*, 2021). Our findings largely corroborated other studies, and this comparison revealed ORF6 as the most consistent and effective inhibitor of IFN induction and ISG signaling across various studies (Fig 1D) (Lei *et al*., 2020), (Li *et al*., 2020), (Xia *et al*., 2020), (Yuen *et al*., 2020), (Zhang *et al*., 2021), (Vazquez *et al*, 2021), (Hayn *et al*., 2021), (Stukalov *et al*., 2021). Hence, we decided to further explore the mechanistic aspects of IFN-antagonism by SARS-CoV-2 ORF6 protein.

**Figure 1.**
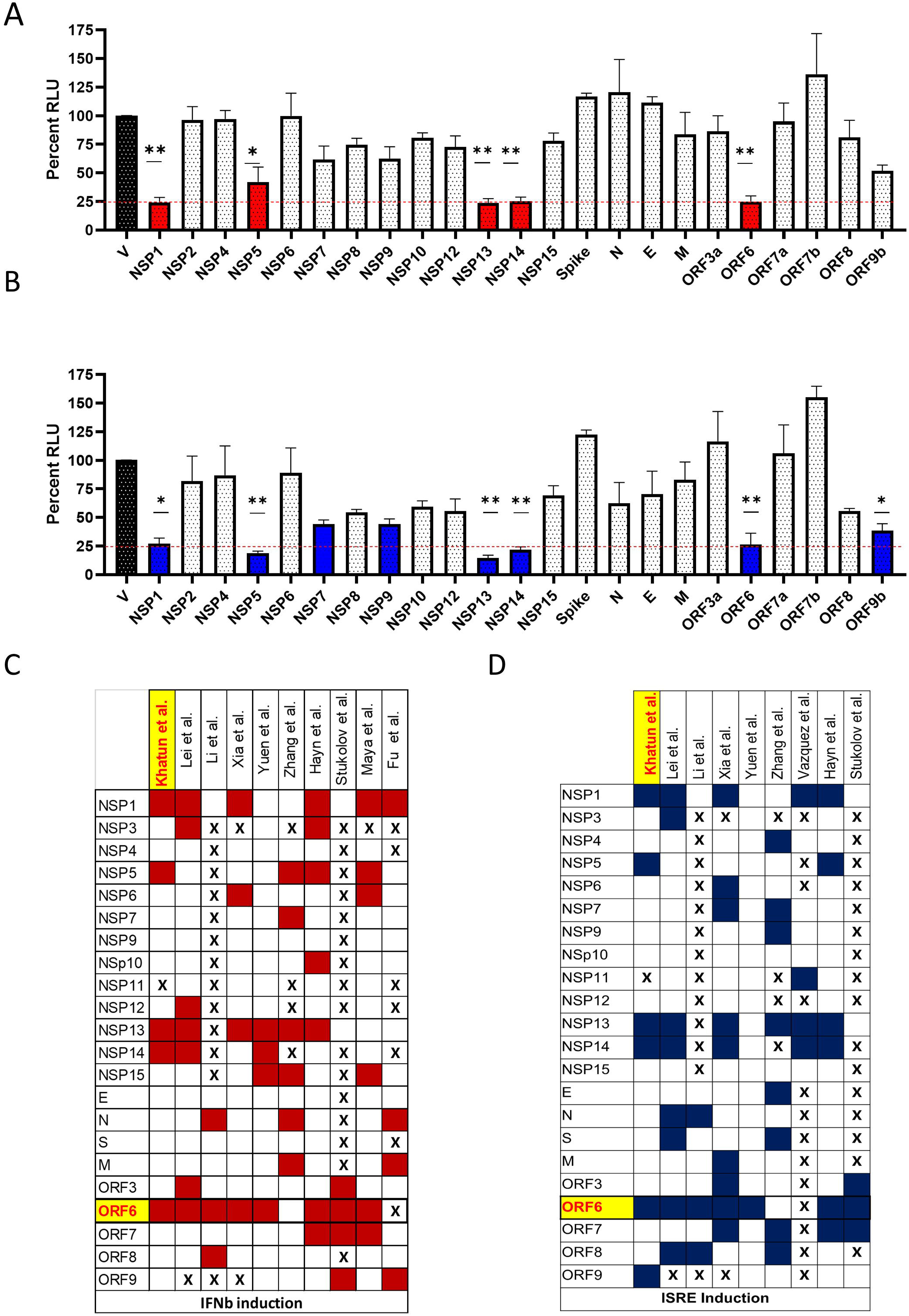
Multiple SARS-CoV-2 proteins antagonize Type-I IFN induction and signaling. A. The graph depicts the quantification of the IFNβ dual luciferase assay. HEK293T cells were co-transfected with IFNβ promoter-driven Firefly luciferase reporter plasmid, Renilla luciferase reporter plasmid, and viral protein-expressing plasmid or empty vector. 24 hours post-transfection, cells were induced with poly I: C, followed by assaying the cells for dual luciferase activity (n=4). B. Quantification of ISRE dual luciferase assay from HEK293T cells which were co-transfected with ISRE promoter-driven Firefly luciferase reporter plasmid, Renilla luciferase reporter plasmid, and viral protein-expressing plasmid or empty vector, followed by induction with Universal type I IFN (1000U/ml) for 12 hours (n=4). C-D. Heatmap of studies that have tested the role of multiple SARS-CoV-2 proteins on IFN induction (C) and ISRE induction (D) pathway using Luciferase assay. Here red and blue color depicts the protein which antagonizes IFN and ISRE induction, respectively. Statistical significance of the data is represented as *P < 0.05; **P < 0.01; ***P < 0.001; ns: not significant. Error bars represent mean + standard error.

### SARS-CoV-2 ORF6 protein inhibits Type-I IFN induction and downstream signaling through distinct mechanisms

To further confirm the IFN-antagonistic activity of SARS-CoV-2 ORF6 protein, we performed dose-response experiments and found that it can inhibit IFNβ and ISRE promoter-driven Luciferase expression in a dose-dependent manner (Sup Fig2A, 2B). In a similar experiment, SARS-CoV-2 ORF6 could inhibit IFNβ-Luciferase induction in Sendai virus (SeV) infected cells (Fig 2A). This was corroborated by measuring the effect of ORF6 expression on IFNβ and ISG54 transcript levels by RT-PCR in SeV-infected cells (Sup Fig 2C). To identify the specific targets of ORF6, we performed an IFNβ Luciferase assay, using different signaling components leading to IFN induction. We observed that ORF6 inhibited IFNβ induced by RIG-I, MAVS, TBK1, IKK□, IRF3-5D, and IRF7-CA to varying degrees, with the most prominent effect seen at the level of RIG-I (Fig 2B-G). This was confirmed in an experiment where ORF6 inhibited RIG-I 2CARD-induced IFNβ reporter activity in a dose-dependent manner (Sup Fig 2D). To further investigate this, we examined the interaction of ORF6 with key mediators of IFN induction and signaling, including RIG-I, MAVS, STAT1, and STAT2, and found that ORF6 interacted with all of them except STAT1 (Fig 2H). Immunofluorescence assays further confirmed this, which showed significant ORF6 colocalization with RIG-I and to a limited extent with MAVS (Fig 2I, Sup Fig 2E). Earlier studies have reported that ORF6 interferes with the nuclear translocation of transcription factors involved in IFN induction (IRF3) and signaling (STAT1). This was corroborated in our experiments where ORF6 expression inhibited SeV infection-induced nuclear translocation of IRF3 and STAT1 (Fig 2J, K). Overall, these data indicated that ORF6 inhibits both type-I IFN and downstream ISG induction by acting upon different cellular components of these pathways.

**Figure 2.**
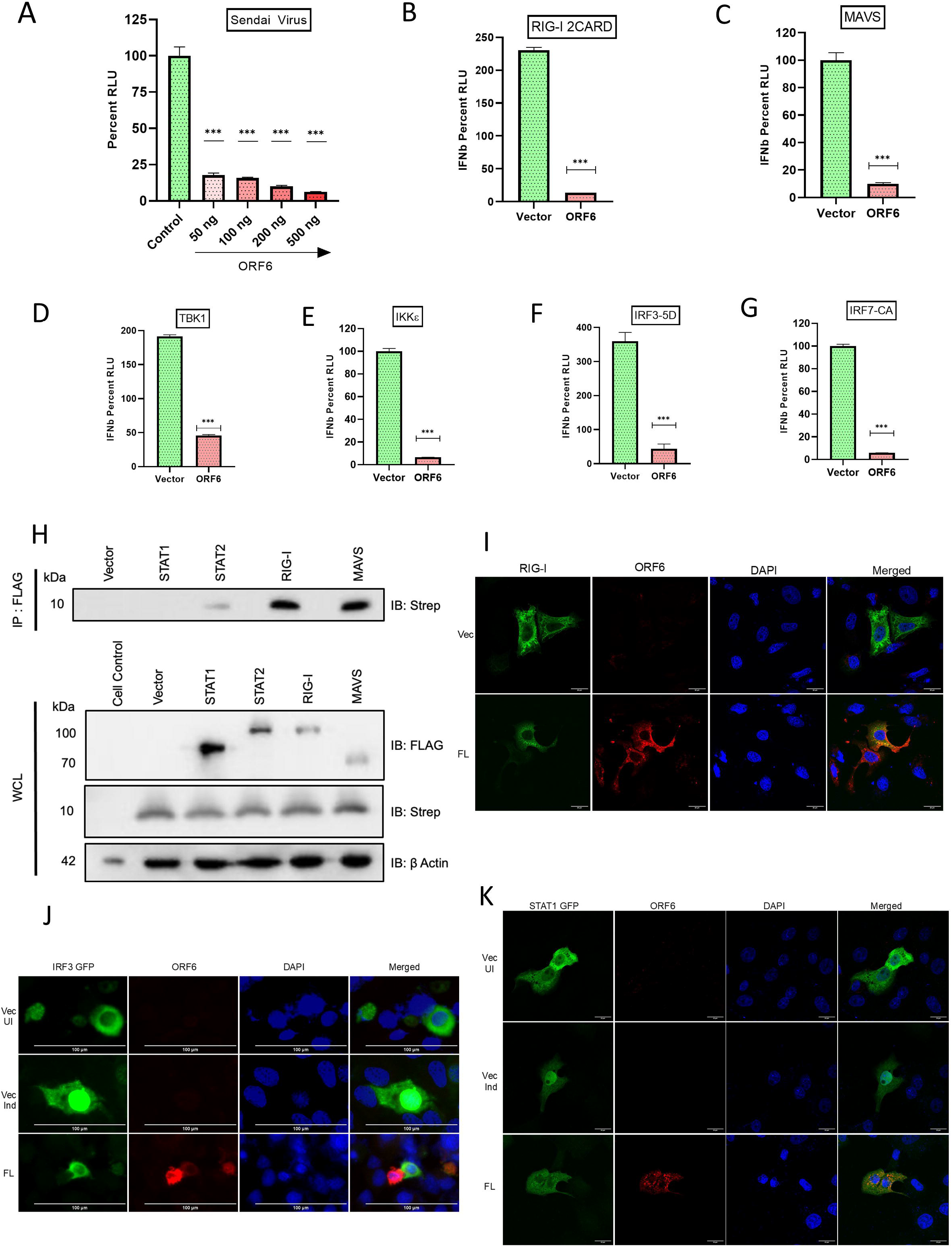
SARS-CoV-2 ORF6 protein inhibits Type-I IFN induction and signaling through distinct mechanisms. **A.** Dual-luciferase assay showing the effect of an increasing amount of ORF6 on IFNβ promoter activation upon Sendai virus infection. (n=4, test details). **B-G**. Dual-luciferase assay depicting the effect of SARS-CoV-2 ORF6 on IFNβ promoter activation in presence of different IFN induction pathway inducer proteins. HEK293T cells were co-transfected with firefly luciferase reporter plasmid driven by IFN β promoter, renilla luciferase reporter, a plasmid expressing ORF6 or empty vector, and a plasmid expressing different proteins of IFN induction pathway. 24 hours -post-transfection cells were lysed for dual-luciferase analysis. H. Coimmunoprecipitation of ORF6-strep and innate immune pathway proteins (Flag-tagged) was performed in HEK 293T cells by overnight incubation of anti-Flag antibody followed by analyzing the eluate by western blot. **I**. Representative confocal images of RIG-I (green) in the presence or absence of ORF6 (red), scale bar 20 μM. **J.** Representative confocal images showing the translocation of IRF3 GFP in the presence or absence of ORF6 (red) upon Sendai virus infection, scale bar 100 μm. **K**. Representative confocal images showing the nuclear translocation of STAT1 GFP in presence of ORF6. Vero E6 cells were co-transfected with ORF6-strep and STAT1 GFP. 24 hours post-transfection, cells were induced with universal IFN (1000 U/ml) for 1 hour, followed by fixation and staining. Scale bar 20 μM. Statistical significance of the data is represented as *P < 0.05; **P < 0.01; ***P < 0.001; ns: not significant. Error bars represent mean + standard error.

### The cytoplasmic domain of SARS-CoV-2 ORF6 is critical for Type I-IFN antagonism

SARS-CoV-2 ORF6 protein is a 61 amino acid long accessory protein (Yuen *et al*., 2020). Its ortholog is present in SARS-CoV but absent in the MERS virus (Chan *et al*, 2020). The C-terminus of the SARS-CoV ORF6 is critical for the innate immune antagonism (Frieman *et al*, 2007). We compared the IFN antagonistic activity of ORF6 proteins from SARS-CoV-2 and SARS-CoV and found them to be similar (Sup Fig 3A). SARS-CoV-2 ORF6 shares a 69% sequence similarity with its SARS CoV counterpart (Yuen *et al*., 2020). Sequence alignment of SARS-CoV-2 with other closely related coronavirus showed that N-terminus cytoplasmic part (M1-Q8) and C-terminus cytoplasmic part (N47-D61) residues are relatively conserved (Sup Fig 3C). This prompted us to map the domains and amino acid residue of SARS-CoV-2 ORF6, essential for type I IFN response antagonism. Structural homology modeling and structure prediction by MA Riojas et al. show the ORF6 protein to be comprised of an N (M1-Q8) and C (N47-D61) terminal cytoplasmic tail and a middle transmembrane domain (V9-E46) (Sup Fig 3B) (Riojas *et al*, 2020). To map the IFN antagonistic activity of ORF6 to its distinct domains, we created expression constructs lacking the N terminus (ΔN), C terminus (ΔC), or expressing only the cytoplasmic tail (C-Cyto) of the protein (Fig 3A). Next, we tested the ability of the deletion construct to inhibit IFN and ISG induction using the Luciferase reporter assay. We observed that ΔC constructs lost the ability to inhibit IFN induction and downstream signaling (Fig 3B, C). When compared, ΔN was still effective in inhibiting IFN induction, but was less effective in restricting ISRE activity than full-length (FL) ORF6 (Fig 3B, C). The importance of the C terminal domain in inhibiting IFN induction was further validated using RIG-I 2CARD and IRF3-5D as inducers; here also ΔC construct was significantly less effective than full-length ORF6 (Fig 3D, E). We also tested the effect of ORF6 domain deletions on RIG-I interaction by co-immunoprecipitation and IRF3 nuclear translocation by immunofluorescence. Although we did not see the loss of interaction with RIG-I with ORF6 upon domain deletion, there was overall reduced expression of RIG-I in the presence of ORF6 (Fig 3F). In western blotting experiments, the C-Cyto construct of ORF6 consisting of only the cytoplasmic tail did not produce a detectable band (Fig 3 F). Furthermore, while the full length and ΔN ORF6 were still effective in restricting IRF3 to the cytoplasm in SeV infected cells, ΔC ORF6 lost this ability. (Fig 3G). Overall, these data established that the C-terminal cytoplasmic domain of the SARS-CoV-2 ORF6 protein is crucial for its ability to restrict both type-I IFN induction and downstream ISG induction.

**Figure 3.**
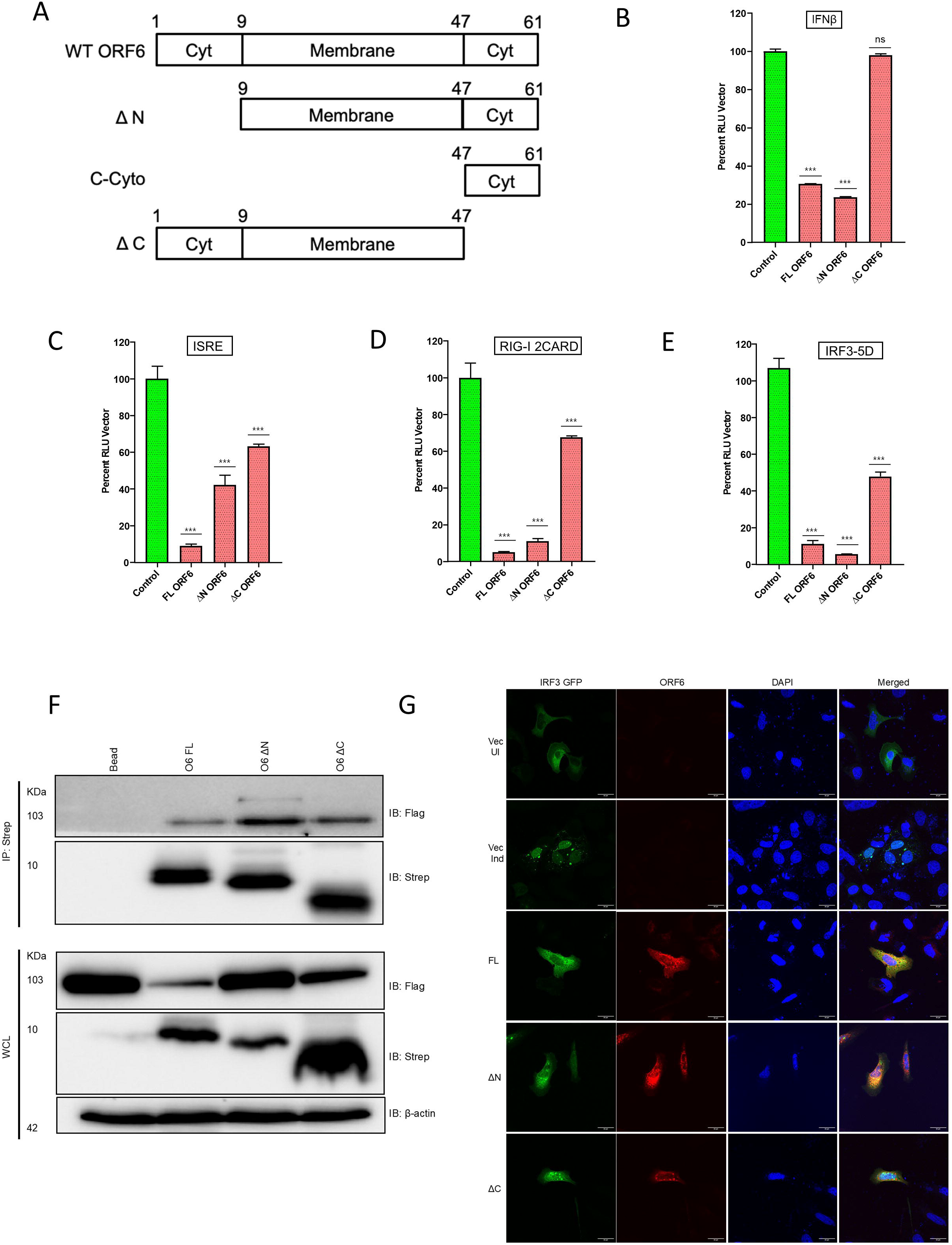
The cytoplasmic domain of SARS-CoV-2 ORF6 is essential for Type I-IFN antagonism. **A.** Schematic representation of the domain structure and involved amino acids of wild type (WT) ORF6, delta N, C-cyto, and delta C. **B-C**. Dual-luciferase assay depicting the effect of full length (FL) and deletions of ORF6 on IFN β (B) and ISRE (C) promoter activation. The principle of the assay is previously described in figure 1 A. **D-E.** Dual-luciferase assay showing the effect of FL and deletions of ORF6 on IFN β promoter activation in presence of an active form of RIG I (RIG I-2CARD) (D) and IRF3 (IRF3-5D) (E). The principle of the assay is previously described in figure 2 B-G. **F.** Coimmunoprecipitation of ORF6-strep (FL and deletions) and RIG-I Flag was performed in HEK 293T cells by overnight incubation of anti-strep antibody followed by analyzing the eluate by western blot. **G**. Representative confocal images performed in HeLa cells showing the translocation of IRF3 GFP upon Sendai virus infection in presence of FL and ORF6 deletions. Scale bar 20 μm. Statistical significance of the data is represented as *P < 0.05; **P < 0.01; ***P < 0.001; ns: not significant. Error bars represent mean + standard error.

### The amino acids 52-61 of SARS-CoV-2 ORF6 protein are essential for its type-I IFN antagonistic activity

The SARS-CoV-2 ORF6 protein has a conserved amino acid stretch from 52-61 aa in the C terminal tail, implicated in IFN antagonism (Fig 4A) (Lei *et al*., 2020). To validate that, we constructed 4 amino acid long alanine scanning mutations with 2 amino acid overlaps, called ORF6 M1 (aa 52-55), M2 (aa 54-57), and M3 (aa 56-59) and M4 (58-61). We tested the expression of these constructs by western blotting, where ORF6 M1 and M2 ran slightly lower compared to the wild-type protein, indicating possible sites of post-translational modification between residues 52 to 57 (Fig 4B). Next, we tested the ability of these ORF6 mutants to inhibit IFN induction due to RIG-I 2CARD and IRF3-5D. The ORF6 M4 showed maximum loss of IFN antagonism against RIG-I 2CARD; however, it behaved like WT ORF6 in the case of IRF3-5D mediated IFN induction (Fig 4C, D). Other mutants from M1 to M3 progressively lost their IFN antagonism against both RIG-I and IRF3. Interestingly, all ORF6 mutants were equally ineffective in mitigating ISRE-driven luciferase expression (Fig 4 E). These data suggest that although the C-terminal tail of ORF6, especially residues 52-56 of ORF6, is crucial for antagonizing IFN induction and signaling, they play distinct roles in antagonizing different components of these signaling processes.

**Figure 4.**
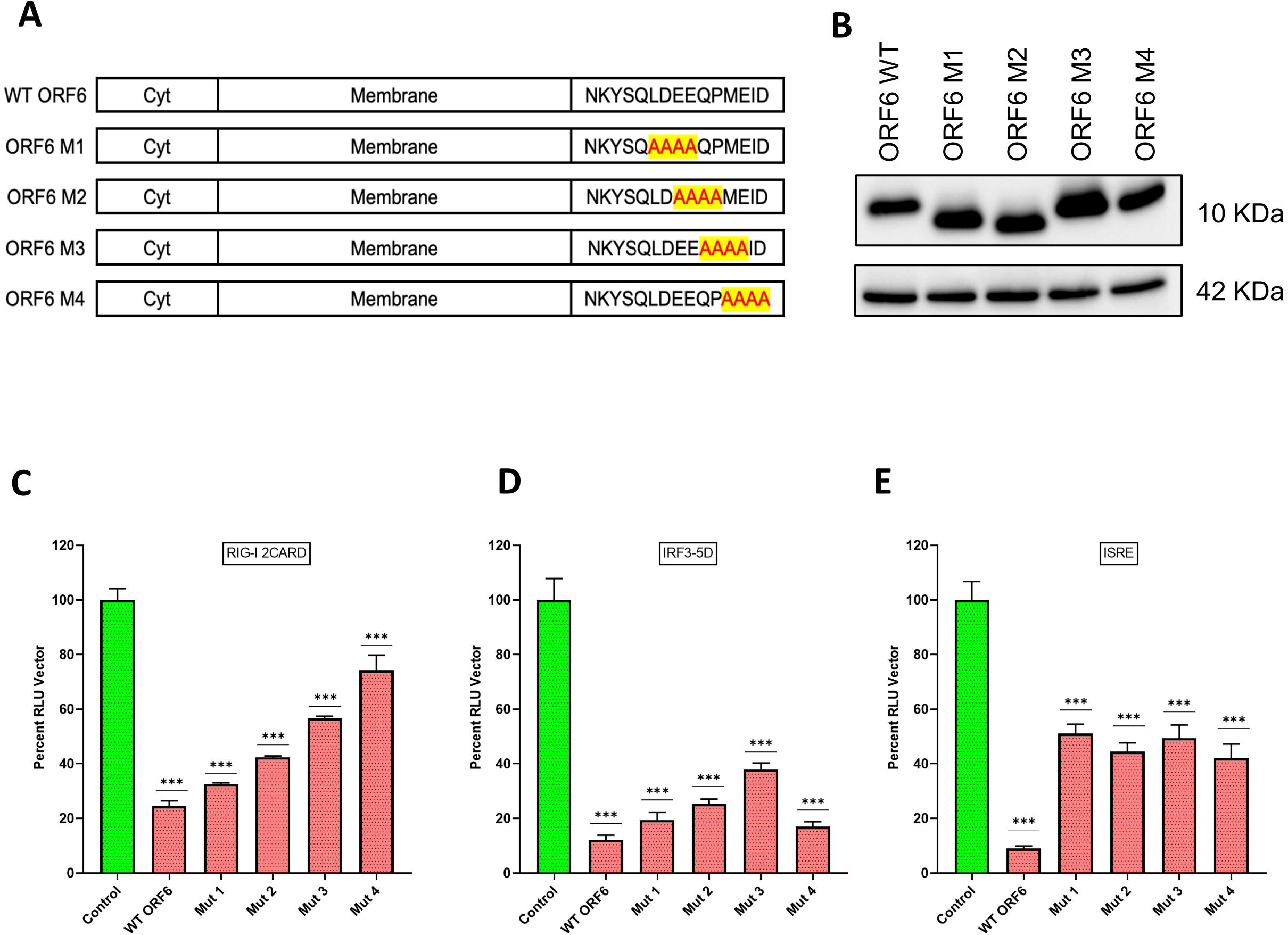
Amino acid residues 52-56 in the C-terminal tail of ORF6 are crucial for its IFN antagonistic function. **A.** Schematic diagram of ORF6 variants. WT: wild type, M1: mutant 1, amino acids 52-55 are substituted with alanine, M2: mutant 2, 54-57 amino acids are substituted with alanine, M3: mutant 3, 56-59 amino acids are converted to alanine, and M4: mutant 4, 58-61 amino acids are converted to alanine. **B.** Western blot analysis with lysates of C-terminally strep-tagged ORF6 variants from HEK293T cells. **C-D**. Quantification of IFN β promoter activation dual luciferase assay of ORF6 variants upon RIG I-2CARD and IRF3-5D induction was performed in HEK293T cells as described previously in figure 2 B-G. **E**. Quantification of ISRE promoter activation of ORF6 variants. The principle of the assay is previously described in Fig 1B. Statistical significance of the data is represented as *P < 0.05; **P < 0.01; ***P < 0.001; ns: not significant. Error bars represent mean + standard error.

### SARS-CoV-2 ORF6 protein targets TRIM25 for degradation to inhibit K63 linked ubiquitination of RIG-I

So far, we have observed that SARS-CoV-2 protein could very potently inhibit RIG-I mediated type-I IFN induction, and its C terminus tail was necessary for this activity. However, deletion of the cytoplasmic tail had no impact on direct interaction between RIG-I and ORF6. Hence, we speculated that perhaps ORF6 interferes with the post-translational modification of RIG-I, which is important for its activity and stability. RIG-I is known to undergo ubiquitination, which can regulate its activity and stability depending on the nature of the linkage (Rehwinkel & Gack, 2020). To this end, we examined wild type, K48 and K63 linked ubiquitination of RIG-I, in the presence or absence of ORF6 in an immunoprecipitation experiment. We observed that the presence of ORF6 reduced overall ubiquitination of RIG-I by wild-type ubiquitin; however, its inhibitory effect was much more prominent against K63-linked ubiquitination than K48 (Fig 5A, B). We also observed reduced expression of RIG-I, in the K63 Ubiquitin overexpression condition in the presence of ORF6 (Fig 5B). Further, we performed a similar experiment to test the ability of SARS-CoV-2 ORF6 deletion constructs to interfere with K63-inked RIG-I ubiquitination. We observed that both full length and ΔN ORF6 were effective; however, ΔC ORF6 lost its ability to reduce K63-linked RIG-I ubiquitination (Fig 5C, D). These data confirmed that ORF6 inhibited K63-linked activating modification of RIG-I to mitigate type I-IFN induction. The K63-linked ubiquitination of RIG-I is mediated by E3 ligase TRIM25 (Gack *et al*, 2007). Thus, we examined the effect of ORF6 on K63-linked ubiquitination of RIG-I in the presence of TRIM25. We observed that not only did TRIM25 mediated K63-linked RIG-I ubiquitination reduce in the presence of ORF6, but the overall TRIM25 expression was also drastically reduced (Fig 6 A, B). This indicated that ORF6 was targeting TRIM25 expression to inhibit its function of RIG-I ubiquitination. Viruses often co-opt cellular proteasome and autophagy machinery to target innate immune signaling mediators for degradation (Choi *et al*, 2018), (Gao & Luo, 2006). To test whether the same applied in the case of ORF6 and TRIM25, we examined the effect of proteasome inhibitor (MG132) and Bafilomycin A1 (BafA) (autophagy inhibitor) on ORF6 mediated reduction of Trim 25 expression. We observed that full-length ORF6 reduced TRIM25 levels, which were partially rescued by both MG132 and BafA (Fig 6C). We also observed that ΔC ORF6 was less effective in reducing TRIM25 expression, which is likely responsible for its loss of type-I IFN antagonism. The proteasomal degradation was also observed in the case of SARS-CoV-2 infected cells and was rescued significantly by MG132 treatment (Fig 6D). Taken together, these data provide a novel mechanism by which SARS-CoV-2 ORF6 inhibits RIG-I mediated type-I IFN induction.

**Figure 5.**
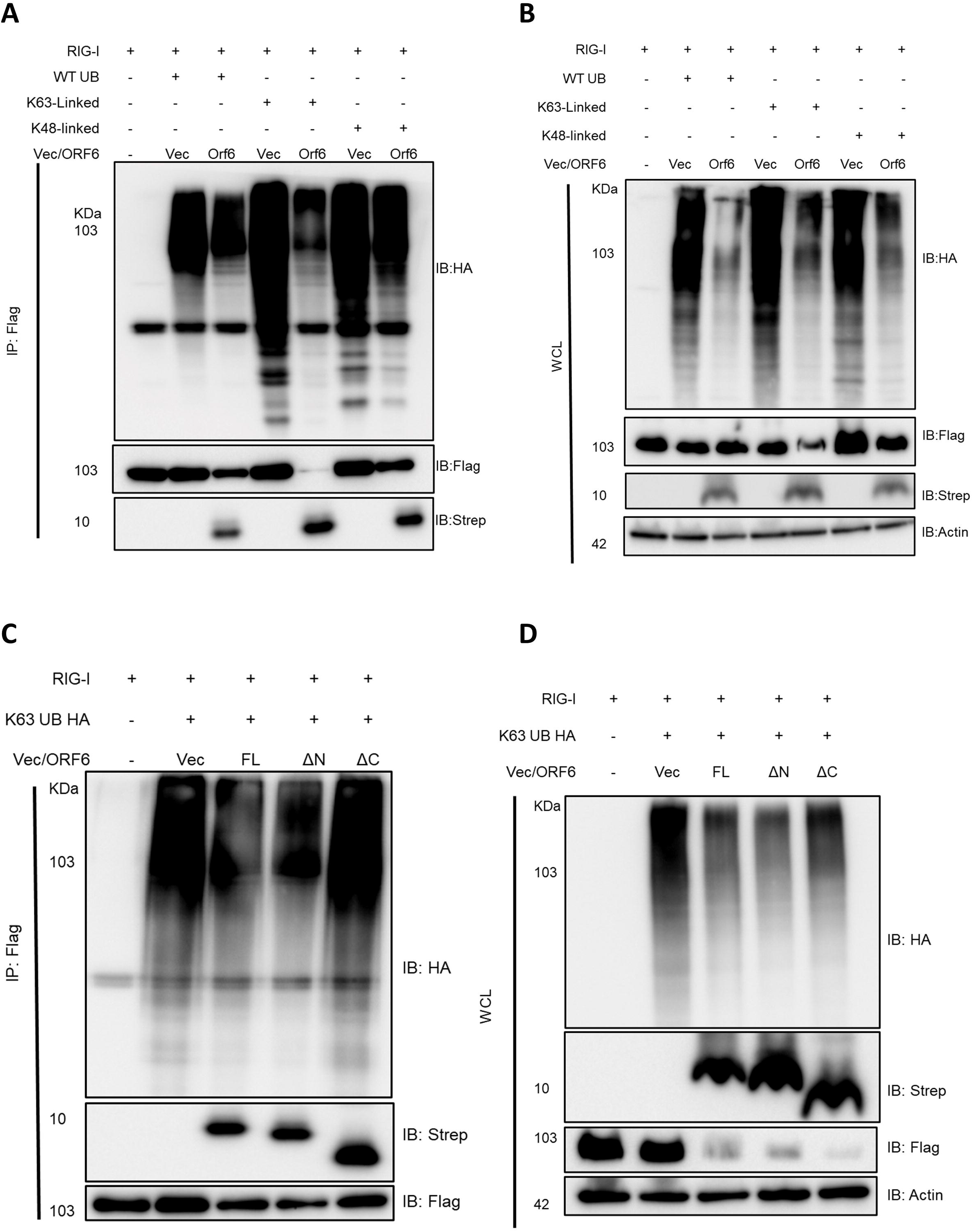
SARS-CoV-2 ORF6 protein Inhibits K63 Linked Ubiquitination of RIG-I. **A-B.** Western blot analysis showing the effect of ORF6-strep on RIG I ubiquitination with cell lysates from HEK293T cells which were co-transfected with RIG I Flag, wild type (WT) UB HA or K63-linked UB HA or K48-linked UB HA and empty vector or ORF6-strep for 48 hours. Cell lysates were either directly analyzed as whole cell lysate (B) or incubated overnight with anti-Flag antibody followed by analysis as IP fraction (A) and probed with indicated antibodies. **C-D**. Western blot analysis with cell lysates from HEK293T cells co-transfected with RIG I Flag, K63-linked UB HA, and different deletions of ORF6-strep. Cell lysates were either directly assessed as WCL (B) or incubated overnight with anti-Flag antibody followed by analysis (A).

**Figure 6.**
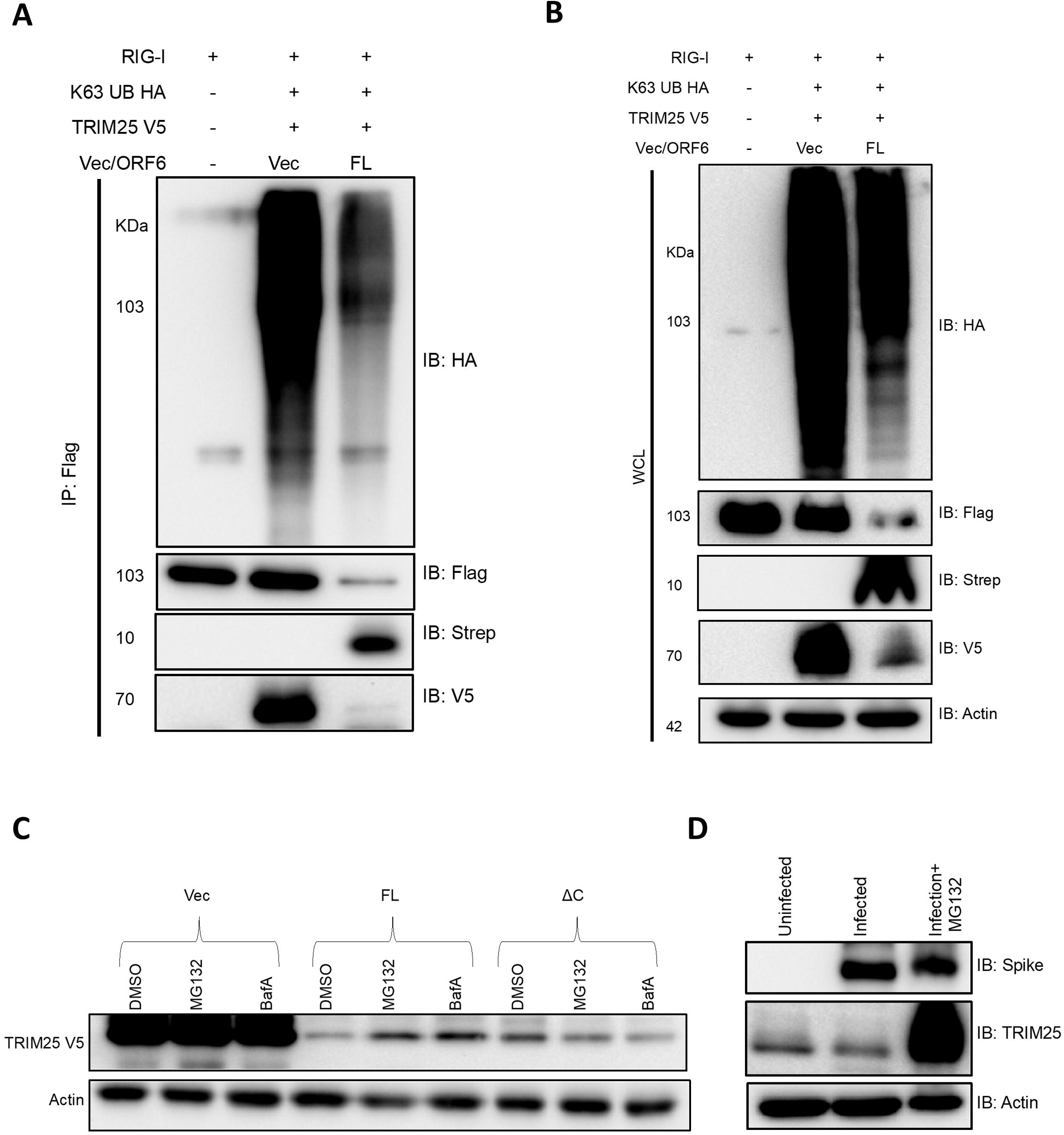
SARS-Cov-2 ORF6 targets Trim25 for proteasomal degradation to inhibit K63-linked Ubiquitination of RIG-I. **A-B.** Western blot analysis with cell lysates from HEK293T cells co-transfected with indicated plasmids. Cell lysates were either directly analyzed as WCL (B) or incubated overnight with anti-Flag antibody followed by eluate analysis by WB and probed with indicated antibodies. **C.** Western blot analysis with cell lysates from HEK 293T cells cotransfected with TRIM25 V5 and strep-tagged ORF6 (FL or ΔC) or empty vector for 36 hours followed by MG132 (5 μM) or BafA (20 nM) treatment for 12 hours. **D.** Western blot analysis with cell lysates from HEK-ACE2 cells infected with SARS-CoV-2. 12 hours post-infection, cells were treated with MG132 (5 μM) for 12 hours, followed by harvesting the cell lysates.

## Discussion

SARS-CoV-2 encodes at least 27 proteins categorized into structural, non-structural, and accessory proteins. While structural proteins like spike are direct targets of humoral immunity and are under constant selection pressure, the accessory and non-structural proteins are often engaged in antagonism and evasion of cellular innate immunity, primarily driven by interferon. The IFN-antagonism by SARS-CoV-2 has been a subject of intense research by several groups, and significant progress has been made in understanding it. In particular, the type-I IFN, which is produced by all respiratory tract epithelial cells to combat viral infection, is targeted by multiple SARS-CoV-2 accessory and non-structural proteins in a myriad of ways, often with overlapping mechanisms (Min *et al*, 2021), (Lei *et al*., 2020), (Li *et al*., 2020), (Xia *et al*., 2020), (Yuen *et al*., 2020), (Zhang *et al*., 2021), (Vazquez *et al*., 2021), (Hayn *et al*., 2021), (Stukalov *et al*., 2021), (Shemesh *et al*., 2021), (Fu *et al*., 2021). This study started with experimentally cataloging the SARS-CoV-2 proteins that either inhibit type-I IFN induction or downstream signaling to produce ISGs or inhibit both. We observed that 4 SARS-CoV-2 proteins (NSP1, NSP13, NSP14, and ORF6) were able to potently inhibit both type-I IFN induction and signaling (Fig 1A, B), which was in accordance with multiple independent studies (Fig 1C, D). The NSP1 protein has been shown to directly interact with ribosomes and inflict general cellular mRNA translation shutdown. This also results in inhibiting the production of IFN and ISGs during viral infection (Thoms *et al*, 2020). The NSP14 protein has been reported to shut down host translation, whereas NSP13 hijacks deubiquitinase USP13 to restrict IFN induction (Hsu *et al*, 2021); (Guo *et al*, 2021). In our experiments, the ORF6 protein was most effective in inhibiting both type-I IFN induction and signaling. Several other research groups have also reported such activity of SARS-CoV-2 ORF6. The mechanism of SARS-CoV-2 ORF6 activity has been mapped to inhibition of nuclear import of crucial transcription factors (STAT1, IRF3) required for IFN response. It does so by associating with the nuclear import co-factor Karyopherin alpha and nuclear pore-component Nup98 (Miorin *et al*, 2020), (Xia *et al*., 2020), (Frieman *et al*., 2007). These interactions also allow ORF6 to restrict the nuclear export of cellular RNAs induced upon infection, which may also contribute to its IFN antagonistic activity (Addetia *et al*, 2021).

In our study, ORF6 was found to exert a strong inhibitory action on RIG-I 2CARD mediated type-I IFN induction, which is a very early step of RLR signaling upstream of nuclear translocation of IRF3 or expression of ISGs. Direct action of ORF6 on RIG-I was not studied before; hence we decided to explore this in more detail. We found that ORF6 directly interacts with RIG-I and leads to its reduced expression. The C-terminal cytoplasmic tail of ORF6 has been reported to be essential for its IFN antagonism (Lei *et al*., 2020). We tested the importance of the same in inhibition of RIG-I function and found that the C-terminal region, especially residues 52-61 of ORF6, were crucial for restricting RIG-I mediated IFN induction, however lack of the C-terminal domain did not affect the direct interaction of ORF6 with RIG-I. This indicated that possibly post-translational modification of RIG-I, which regulates its activity and stability, may be affected by ORF6. Upon sensing a viral PAMP, the RIG-I protein undergoes the K63 linked ubiquitination by E3 ligase TRIM25, essential for its activation and downstream signaling (Gack *et al*., 2007). This step of RLR signaling is often targeted by viral proteins, especially RNA viruses (Liu *et al*, 2016). Our study showed that TRIM25 mediated K63-linked ubiquitination of RIG-I was significantly reduced in the presence of ORF6. This inhibitory effect was partially lost upon deletion of the C-terminal domain of ORF6, which explains the requirement of this region for type-I IFN antagonism. Further exploration of the effect of ORF6 on TRIM25 and RIG-I revealed that ORF6 targets TRIM25 for degradation. This phenotype is reversed partially by proteasome and autophagy inhibitors as well as by deletion of the C-terminal domain of ORF6. The degradation of TRIM25 was also observed in the case of SARS-CoV-2 infection, which was reversed upon treatment with a proteasomal inhibitor. RIG-I and its TRIM25 mediated ubiquitination have also been reported to be targeted by the NSP5 and nucleoprotein (N) of SARS-CoV-2 (Wu *et al*, 2020b); (Zhao *et al*, 2021). An additional layer of inhibition of the same by ORF6 highlights the significance of RIG-I signaling during SARS-CoV-2 infection.

Based on the collective results obtained in our study, we have proposed a working model for SARS-CoV-2 ORF6 mediated inhibition of type I-IFN induction, shown in Figure 7. How ORF6 targets TRIM25 for degradation remains to be investigated and can potentially present an avenue for therapeutic intervention against SARS-CoV-2. ORF6 is a relatively conserved protein of SARS-CoV-2; however, few natural amino acid deletions have been observed in this protein (Queromes *et al*, 2021), (Riojas *et al*., 2020), and it will be interesting to examine the role of the same on the IFN-antagonism function of ORF6. Finally, ORF6 and other IFN antagonists of SARS-CoV-2, which may be dispensable for viral replication, can be removed from the viral genome to engineer attenuated strains, potentially useful as vaccines.

**Figure 7.**
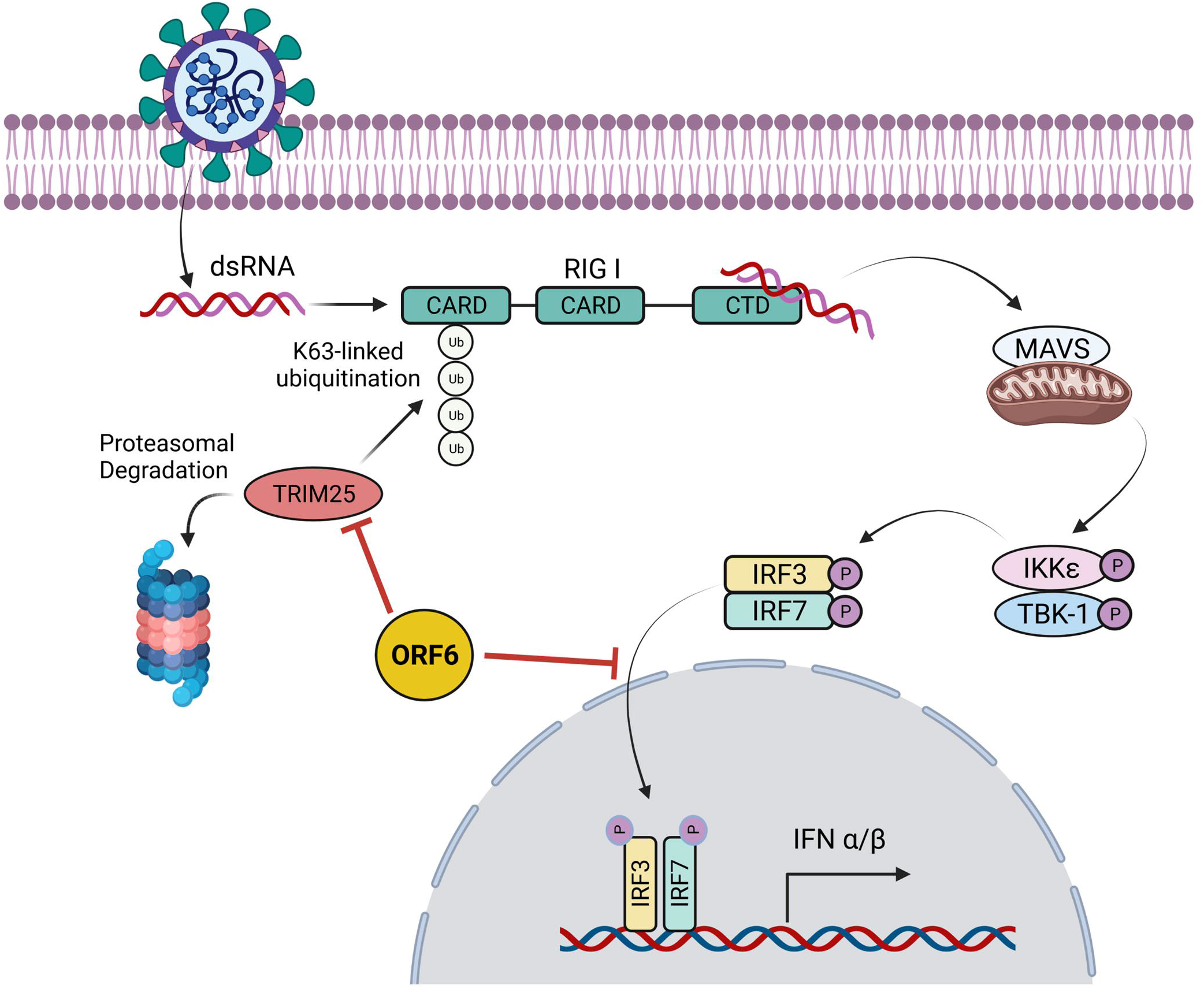
Model of SARS-CoV-2 ORF6 mediated inhibition of K63-linked Ubiquitination of RIG-I. Pictorial representation of stages of IFN induction pathway upon SARS-CoV-2 RNA recognition where TRIM25 mediates K63-linked ubiquitination of RIG-I CARD domain, which in turn gets activated and activates the downstream signaling pathway. In this study, we show that SARS-CoV-2 ORF6 inhibits nuclear translocation of IRF3 and targets TRIM25 for proteasomal degradation, which results in down-regulation of K63-linked ubiquitination of RIG-I.

## Materials & Methods

### Plasmids & Antibodies

Plasmids expressing SARS-CoV-2 proteins were a kind gift from Prof. Nevan Krogan (University of California San Francisco) and have been described before (Gordon *et al*, 2020). The IFN induction and signaling plasmids (IFN□-Firefly Luciferase, ISRE-Firefly Luciferase, RIG-I, 2-CARD, TBK1, IKK□, IRF3-5D, IRF7-CA, STAT1, STAT2) were provided by Prof. Adolfo García-Sastre (Icahn School of Medicine at Mount Sinai, New York) and have been described before (Versteeg *et al*, 2013). For constructing the deletion mutants, the desired sequence was PCR amplified from pLVX-EF1alpha-SARS-CoV-2-ORF6-2xStrep-IRES-Puro plasmid, followed by cloning in the pCAGGS backbone using EcoRI (ER0271, Thermo scientific) and XhoI (ER0691, Thermo scientific) restriction enzyme including the full-length ORF6. Delta C plasmid was constructed using overlap extension PCR. ORF6 variant mutants were constructed by subcloning the construct in a TA backbone, followed by substituting the residues as described before (Edelheit *et al*, 2009) and cloning in the pCAGGS backbone using EcoRI and XhoI.

### Cell Lines and cell culture

Human embryonic kidney 293T (HEK293T), A549 lung adenocarcinoma, and HeLa cells were purchased from National Centre for Cell Science (NCCS), and HEK-ACE2, Vero cells were procured from the America Type Culture Collection (ATCC, Bethesda, MD). Cells were cultured in complete media containing high-glucose Dulbecco’s modified Eagle’s medium (DMEM) (12100-046, GIBCO) with 10% FBS (16140-071, Gibco), 100 IU/ml Penicillin, 100 mg/ml Streptomycin and 0.25mg/ml, supplemented with GlutaMAX^™^ (35050-061, Gibco) at 37°C with 5% CO2.

### Viruses and infection

#### Infection

SARS-CoV2 (Isolate Hong Kong/VM20001061/2020, NR-52282, BEI Resources, NIAID, NIH) was propagated and titered by plaque assay in Vero E6 cells as described before (Case *et al*, 2020).

HEK-ACE2 (for SARS-CoV-2) or HEK293T (SeV) were seeded in a 24-well cell culture dish (pre-coated with 0.1 mg/mL poly-L-lysine (P9155-5MG, Sigma-Aldrich) for HEK-ACE2) and 24 hr later, used for infection. Infection was done with 2 MOI SARS CoV-2 in DMEM supplemented with 10% FBS or 100 HAU SeV in OPTI-MEM in 100 μl inoculum for 1hr at 37°C. Complete DMEM was added to the cells.

#### Plasmid transfection

HEK-293T cells (0.1 X 10^6^ cells/well) were seeded in a 24 well plate pre-coated with 0.1 mg/mL poly-L-lysine (P9155-5MG, Sigma-Aldrich) and 24 hours later used for transfection. Cells were then transfected with 0.5μg of expression plasmid using lipofectamine-2000 reagent (11668019, Invitrogen) or lipofectamine-3000 (L3000015, Invitrogen), according to the manufacturer’s instruction.

### Luciferase Reporter Assay

For the IFN induction assay, HEK-293T cells were co-transfected, in duplicates, with 50 ng of IFNβ-luc firefly luciferase reporter plasmid, and 20 ng of pRL-TK Renilla luciferase reporter plasmid along with 500 ng of SARS-CoV-2 protein expression plasmid or empty vector. 24 hours post-transfection, cells were induced with poly (I: C) (1μg), or 100 HAU of Sendai virus for 12 hours, followed by lysing of the cells for analyzing the dual-luciferase activity. For dissecting the steps of the IFN induction pathway, HEK-293T cells were co-transfected with 50 ng of inducer plasmid along with the above-indicated plasmids. 24 hours post-transfection, cells were lysed. Similarly, for IFN signaling assay, HEK-293T cells (0.1 X 10^6^ cells/well in a 24 well plate) were co-transfected with 50 ng of ISRE-luc firefly luciferase reporter plasmid and 20 ng of pRL-TK renilla luciferase reporter plasmid along with 500 ng of SARS-CoV-2 protein expression plasmid. 24 hours post-transfection, the cells were induced with 1000U/ml of Universal Type I IFN (PBL assay, Cat No. 11200-1). 12 hours post-induction, the cells were lysed, and luciferase activity was measured using a dual-luciferase assay system (E1980, Promega) according to the manufacturer’s instructions. Firefly and Renilla luciferase signals were quantified using Tecan plate reader (INFINITE M PLEX). The signals were represented as percentage fold change with respect to the induced vector.

### Immunofluorescence Assays

A549 cells were seeded on coverslips in a 24-well plate (0.1 X 10^6^ cells per well) for overnight incubation. They were then co-transfected with 500 ng of IRF3-GFP and 500 ng of SARS-CoV-2 protein-expressing plasmid using Lipofectamine 2000 reagent (11668019, Invitrogen). 24 hours post-transfection, the cells were induced with 6 hours of Sendai virus infection (500 HAU). Similarly, Vero cells were seeded on coverslips in 24 well plates (0.1 X 10^6^ cells/well) for overnight incubation. Cells were then transfected with 250 ng of STAT1-GFP along with 250 ng of SARS-CoV-2 viral protein-expressing plasmids. 24 hours post-transfection, cells were induced for 1 hour with 1000U/ml universal Type I interferon (IFN) (11200-2, PBL Assay Science). Similarly, HeLa cells were transfected with 250 ng of RIG I Flag or MAVS Flag and 250 ng of ORF6 strep for 24 hours. Cells were washed with 1X PBS 162528, MP Biomedicals) twice at room temperature and fixed in PBS with 4% formaldehyde (Q24005, Qualigen) at room temperature for 20 min. The cells were then permeabilized using 1% Tween-20 (P1379, Sigma-Aldrich) in PBS at room temperature for 10 min. After three washes, cells were treated with 2% BSA (0215240105, MP Biomedicals) blocking at room temperature for 2 hours, followed by overnight incubation with primary antibody at 4°C and then 2 hours of secondary antibody incubation at room temperature after three washes. The cells were then counter-stained with 4’,6-diamidino-2-phenylindole (DAPI), (D9542-10MG, Sigma-Aldrich) for 10 minutes at RT. Coverslips were mounted on a slide using antifade mounting media (P36970, Invitrogen) and imaged using Zeiss 880.

### Co-immunoprecipitation and Immunoblotting

Transfected cells were washed with ice-cold 1X PBS and then were lysed using 500 *μ*l Pierce lysis buffer per 100 mm dish. The lysate was incubated on ice for 30 minutes with vigorous vortexing every 10 minutes. Samples were sonicated (25% Amplitude, 5 sec ON – 5 sec OFF for 2 cycles) and clarified by centrifugation at 13,000 rpm for 10 min at 4°C. The clarified supernatant was either stored at –80°C or analyzed by immunoblotting as whole cell lysate (Input). Immunoprecipitation was performed for the remaining lysates by overnight incubation with specific antibodies. Subsequently, the complex was pulled down by magnetic beads coated with protein-G (88847, Invitrogen) according to the manufacturer’s instructions. Elution was done directly by 1X Laemmli buffer (1610747, BIO-RAD). Protein samples were resolved by SDS-polyacrylamide gel electrophoresis, followed by transfer onto a PVDF membrane (IPVH00010, Immobilon-P; Merck). Blocking was performed at room temperature for 2 hours using 5% skimmed milk (70166, Sigma-Aldrich) in 1X PBS containing 0.05% Tween 20 (P1379, Sigma-Aldrich) (1X PBST) (or 5% BSA (0215240105, MP Biomedicals) for phospho-proteins). Required primary antibody incubation was performed overnight (12 hr) at 4°C with slow rocking. Secondary antibody incubation was performed at room temperature for 2 hours using Goat Anti-Mouse IgG H&L (31430, Thermo) or Goat Anti-Rabbit IgG H&L (31460, Thermo). The proteins were visualized using Clarity Western ECL Substrate (1705061, BIO-RAD).

### Quantitative Real-Time PCR

1 μg of RNA was reverse transcribed into cDNA using Prime Script^™^ RT Reagent Kit with gDNA Eraser (Perfect Real Time) (RR047A, Takara-Bio) and then diluted 5-fold with nuclease-free water. The gene expression study was conducted using PowerUp™ SYBR^™^ Green Master Mix (A25778, Applied Biosystems^™^) with 18S rRNA as the internal control and appropriate primers for the genes.

### Graphical representations and statistical analysis

All numerical data of luciferase assays and qRt-PCR were analysed and plotted using GraphPad Prism v8.0.2. Statistical significance was calculated using t-test with Bonferroni corrections for multiple comparisons wherever necessary). The P values were indicated as *P < 0.05; **P < 0.01; ***P < 0.001; ns = not significant. Error bars represent mean + standard error. The model diagram of ORF6 action (Fig 7) and 3-D structure (Sup Fig 3C) were made by using Biorender

## Acknowledgments

This work is supported by funding from DBT-Wellcome Trust India Alliance to ST Lab. We acknowledge Infrastructure support from the DBT-IISc partnership, DBT-BIRAC, Crypto Relief Fund, L & T Trust, and DST-FIST program to IISc. We are grateful to Prof. Adolfo Garcia Sastre, Microbiology Department, Icahn School of Medicine, New York, and Prof. Nevan J. Krogan, Cellular Molecular Pharmacology, University of California, San Francisco, for providing plasmids for studying IFN response and SARS-CoV2 protein expression respectively. We thank Rajesh Thangavel Yadav for the help with creating 3D strutre model of ORF6.

## Author Contributions

ST conceived the study, designed the experiments, analyzed the data, and wrote the draft. OK, MS and RN performed the experiments, analyzed the data, and revised the draft.

## Data and Reagent Availability

All data are included in the manuscript. Requests for reagents can be sent by email to Shashank Tripathi.

---

**Supplementary Figure 1. Expression of SARS-CoV-2 proteins and quantification of their inhibition of Type-I IFN induction and signaling.**

A. Western blot analysis with lysates of C-terminally strep-tagged SARS-CoV-2 proteins from HEK293T cells. The position of the respective protein is shown with a red arrow. Non-expressive proteins are shown inside the red square. B. Heatmap representing the percentage RLU and corresponding p-value of each protein from IFNβ and ISRE luciferase screening assay. Here green is >100%, yellow is <50%, pink is <30%. Significant p values are shown in red color.

**Supplementary Figure 2. SARS-CoV-2 ORF6 inhibits Type-I IFN induction and signaling through distinct mechanisms.**

**A, B, and D**. Dual-luciferase assay performed in HEK293Tcells showing the effect of an increasing amount of ORF6 on (A & D) IFN β promoter activation in presence of different inducers like poly I: C (A), RIG I-1CARD (D) and (B) effect on IFN signaling pathway upon universal IFN induction. (test details). **C.** mRNA levels of IFN β induction pathway protein (IFNb) and signaling pathway protein (ISG54) were measured from total RNA from HEK293T cells transfected with ORF6 or empty vector followed by Sendai virus infection. (test details). **E.** Representative confocal images of MAVS (green) in the presence or absence of ORF6 (red), scale bar 20 μM. Statistical significance of the data is represented as *P < 0.05; **P < 0.01; ***P < 0.001; ns: not significant. Error bars represent mean + standard error.

**Supplementary Figure 3. Sequence conservation and predicted three-dimensional structure of ORF6**

**A.** The graph depicting the comparison of IFN β promoter activation in HEK293T cells upon poly I: C induction between SARS-CoV and SARS-CoV-2 ORF6. (test details). **B.** Three-dimensional model structure prediction using I-TASSER (Iterative Threading Assembly Refinement) standalone software version 5, which was visualized using Jmol 14.32. The final images were made using Biorender. **C.** Multiple sequence alignment of ORF6 amino acid sequence from indicated coronaviruses (top). ≠ Indicates conserved amino acids. Schematic representations of cytoplasmic (Cyt) and membrane domains of ORF6 are indicated at the bottom, along with involved amino acids. Statistical significance of the data is represented as *P < 0.05; **P < 0.01; ***P < 0.001; ns: not significant. Error bars represent mean + standard error.

## Tables & Their Legends

| Antibodies | Source | Identifier |
| --- | --- | --- |
| Anti-Strep-tag II antibody | abcam | ab76949 |
| Monoclonal ANTI-FLAG® M2 antibody | Sigma-Aldrich | F3165-.2MG |
| DYKDDDDK Tag (D6W5B)<br>Rabbit mAb(Binds to same epitope as Sigma's Anti-FLAG® M2 Antibody) | Cell Signalling Technologies | 14793S |
| Beta Actin Monoclonal Antibody (BA3R), HRP | ThermoFisher Scientific | MA5-15739-HRP |
| Biotin Anti-HA tag antibody | abcam | ab26228 |
| V5-Tag (D3H8Q) Rabbit mAb | Cell Signalling Technologies | 13202 |
| TRIM25 Monoclonal Antibody (5B5B12) | ThermoFisher | MA5-31935 |
| SARS-CoV / SARS-CoV-2 (COVID-19) spike antibody | GeneTex | GTX632604 |
| Goat Anti-Mouse IgG H&L | ThermoFisher | 31430 |
| Goat Anti-Rabbit IgG H&L | ThermoFisher | 31460 |

