## Supplementary figures and images for "ORF6 protein of SARS-CoV-2 inhibits TRIM25 mediated RIG-I ubiquitination to mitigate type I IFN induction"

### Sup Figure 1

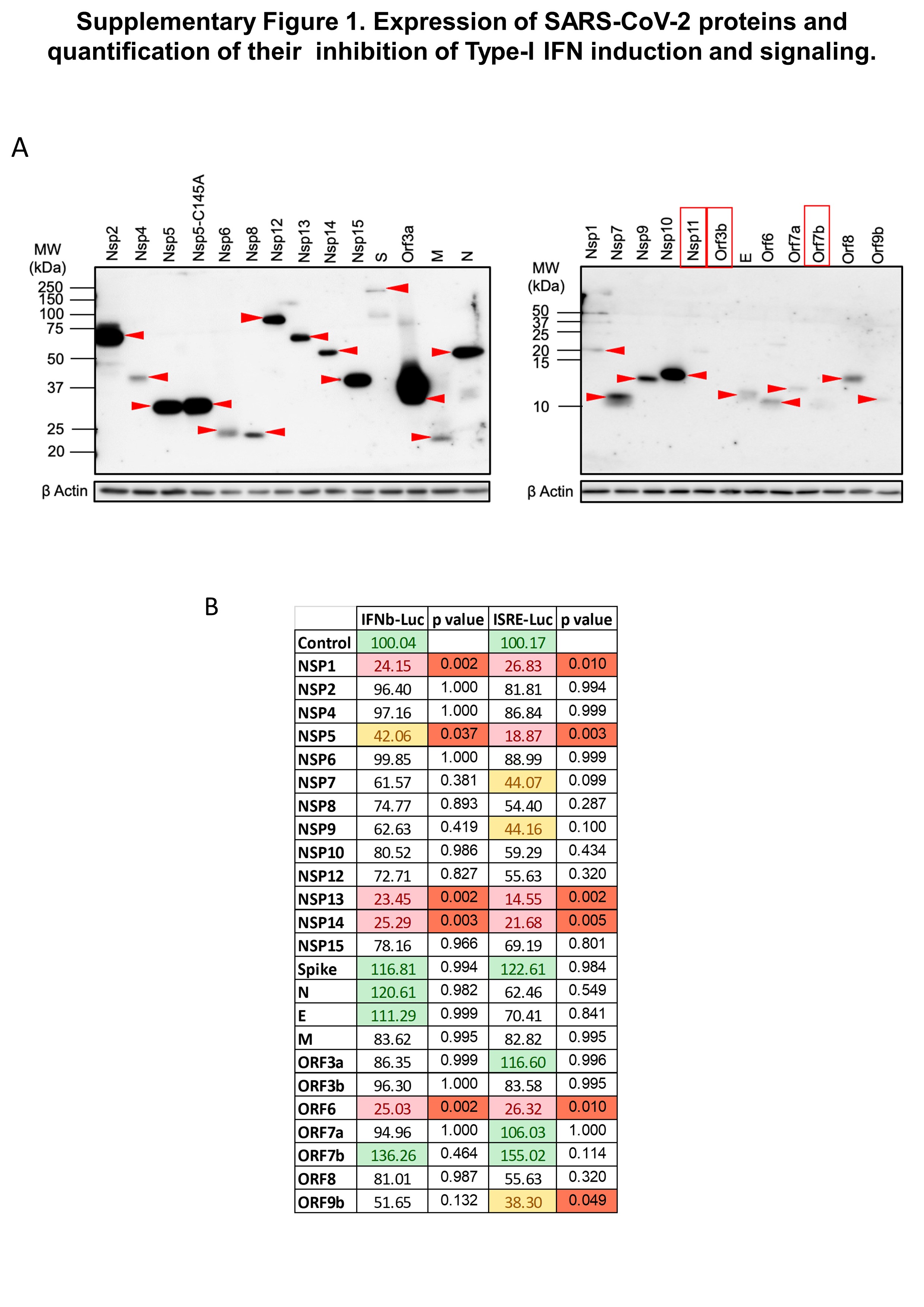

### Sup Figure 2

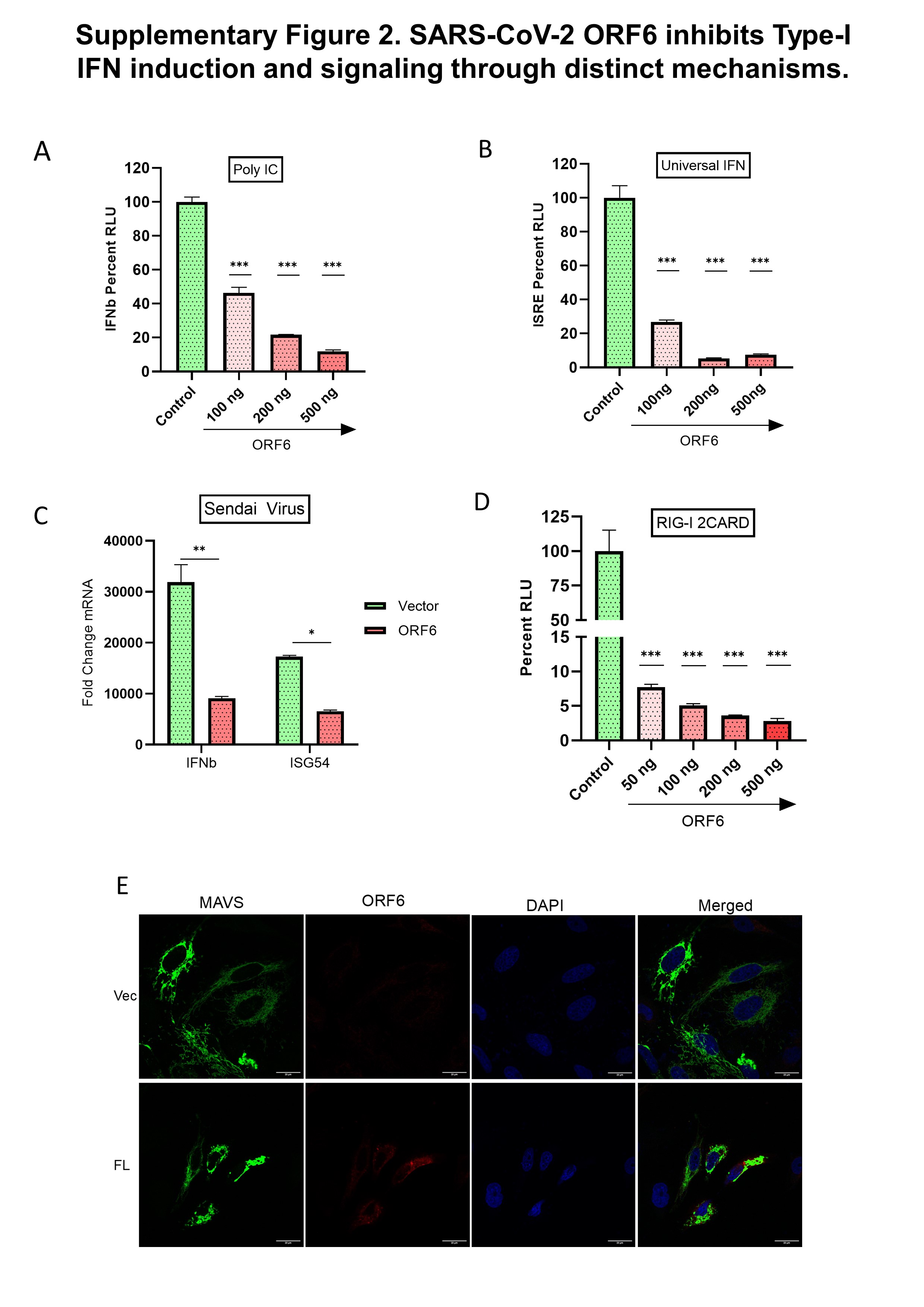

### Sup Figure 3

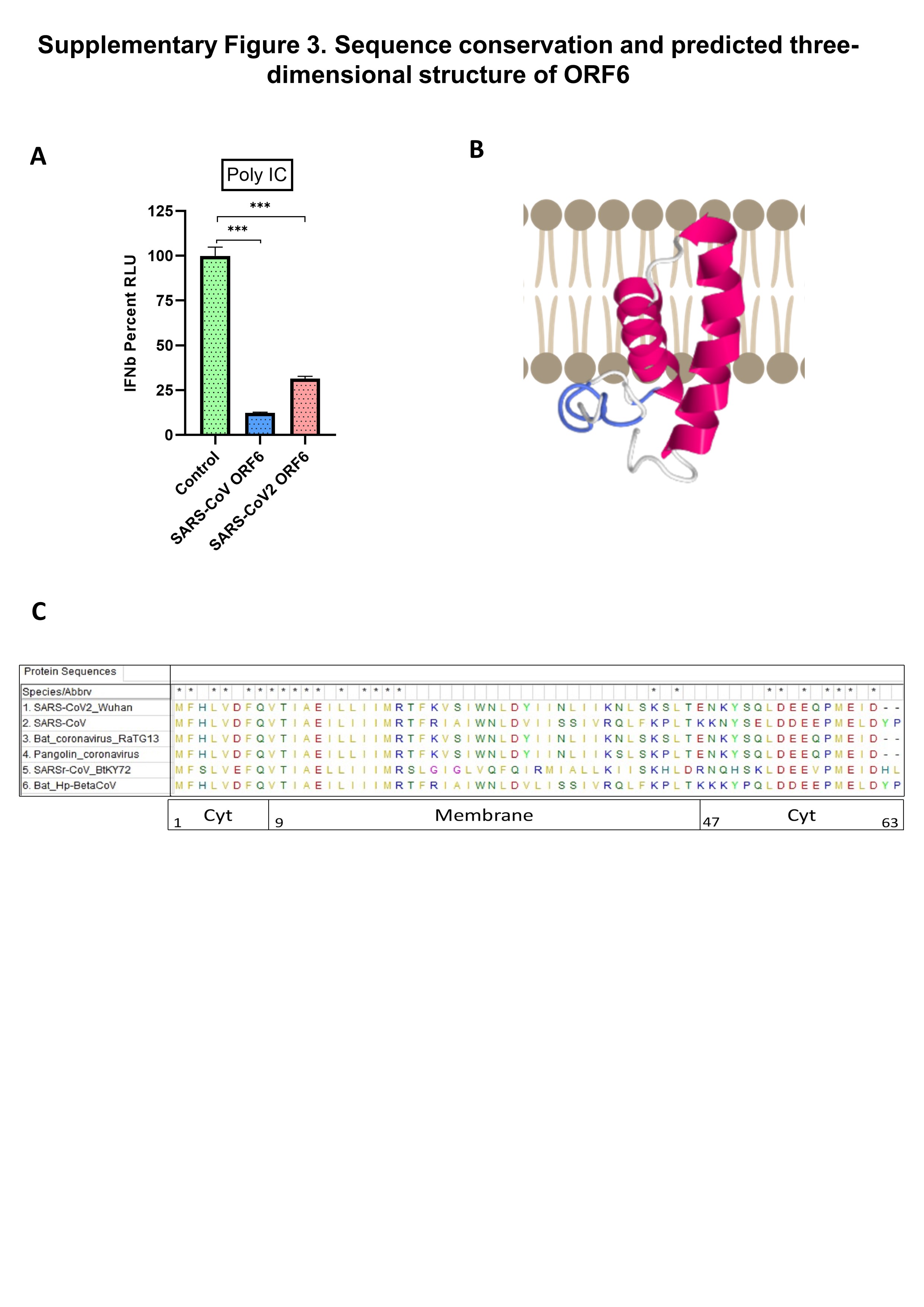
